# Tailocins are recurrent contaminants in ex vivo amyloid extraction

**DOI:** 10.64898/2026.06.17.733002

**Authors:** Jan-Hannes Schäfer, Robert T. O’Neill, Zi Gao, Danielle Grotjahn, Evan T. Powers, Jeffery W. Kelly, Gabriel C. Lander

**Affiliations:** Department of Integrative Structural and Computational Biology, Scripps Research; La Jolla, CA, USA; Department of Chemistry, Chi-Huey Wong Laboratories for Biomedical Research, Scripps Research; La Jolla, CA, USA

**Keywords:** contamination, tailocin, helical reconstruction, amyloidosis, cryoSPARC

## Abstract

Amyloid fibrils extracted from patient tissue are enriched by physico-chemical fractionation methods and enzymatic tissue digestion with collagenase, resulting in a mixture of sedimented complexes. Reprocessing of an ex vivo lysozyme amyloid dataset revealed non-amyloid tubular assemblies, which we reconstructed to 3.5 Å. Sequence-agnostic model building with CryoAtom and fold search with FoldSeek identified a phage tail tube fold. Testing the crude Clostridium histolyticum collagenase used for tissue digestion, negative-stain TEM and LC-MS/MS showed proteins of an F-type tailocin, allowing us to assign the helical reconstruction to a major tail tube protein (TTP) of C. histolyticum. We also observed tailocin contamination in cardiac light-chain amyloid extracts, following the same collagenase treatment. Overall, we identify tailocins as recurrent reagent-borne contaminants in collagenase-treated ex vivo amyloid preparations and propose strategies to detect and avoid such artefacts in single-particle cryo-EM by optimizing the experimental design and data annotation towards a database of common contaminants.

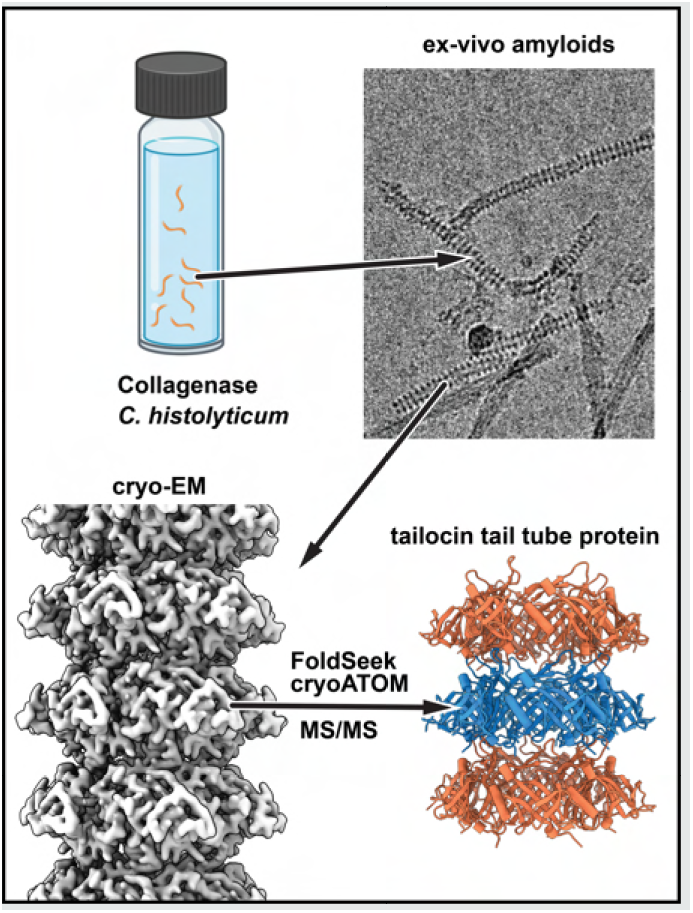

## Introduction

Extraction of disease-relevant amyloid fibrils from patient tissue has become the standard for cryo-electron microscopy studies, linking fibril morphology to a specific pathology ^24,36^. While amyloid fibrils from patients with amyloidosis of the central nervous system can generally only be obtained from postmortem specimens, systemic amyloids can be extracted from living patients following organ transplants, operative biopsy, or lately punch-biopsies ^30,46^. Transthyretin amyloidosis (ATTR) and Light-Chain amyloidosis (AL) manifest across several organ systems, allowing patient-derived structures to now span hereditary and wild-type disease, multiple organs from single individuals, and different polymorphs ^17,29,37,42^. Established amyloid extraction protocols utilize tissue digestion with collagenase, a step introduced by Annamalai and colleagues ^2^ as a modification of the classical Pras protocol ^39^ and subsequently adopted throughout the field ^13,29,37,41,42^. Extraction protocols traditionally use collagenase from Clostridium histolyticum culture filtrates rather than purified enzyme preparations, on the assumption that non-collagenolytic components are either inert or removed during downstream sample preparation. Re-processing cryo-EM datasets can reveal unexpected findings, as demonstrated by the identification of vault particles in ex vivo tau samples ^27^. Similarly, this approach proved informative for deposited datasets of lysozyme amyloids associated with systemic ALys amyloidosis. Here, we present the helical reconstruction of the major tail tube protein (TTP) of a tailocin from Clostridium histolyticum, which was co-purified with amyloid fibrils extracted from patient tissue affected by lysozyme amyloidosis. Using the same collagenase treatment and extraction approach, we identified tailocins in cardiac amyloid extracts from a patient with light-chain amyloidosis, showing that the same reagent-derived contaminant can recur across distinct ex vivo amyloid preparations that use Clostridium histolyticum collagenase.

## Results & Discussion

### Structure of a *Clostridium histolyticum* tailocin

During re-processing of an ex vivo lysozyme variant D87G ^21^ dataset (EMPIAR-11785), associated with ALys, we identified non-amyloid fibrillar assemblies by 2D classification (**Fig. 1A**). Approximately one-third of micrographs contained these distinct fibrillar structures. Helical refinement using a rise of 37.23 Å and twist of 14.17° (**Fig. S1**) yielded a 3.5 resolution reconstruction of 10 nm diameter tubes with a C6-symmetric protofilament arrangement and an inner diameter of 3.5 nm (**Fig. 1B**).

**Figure 1.**
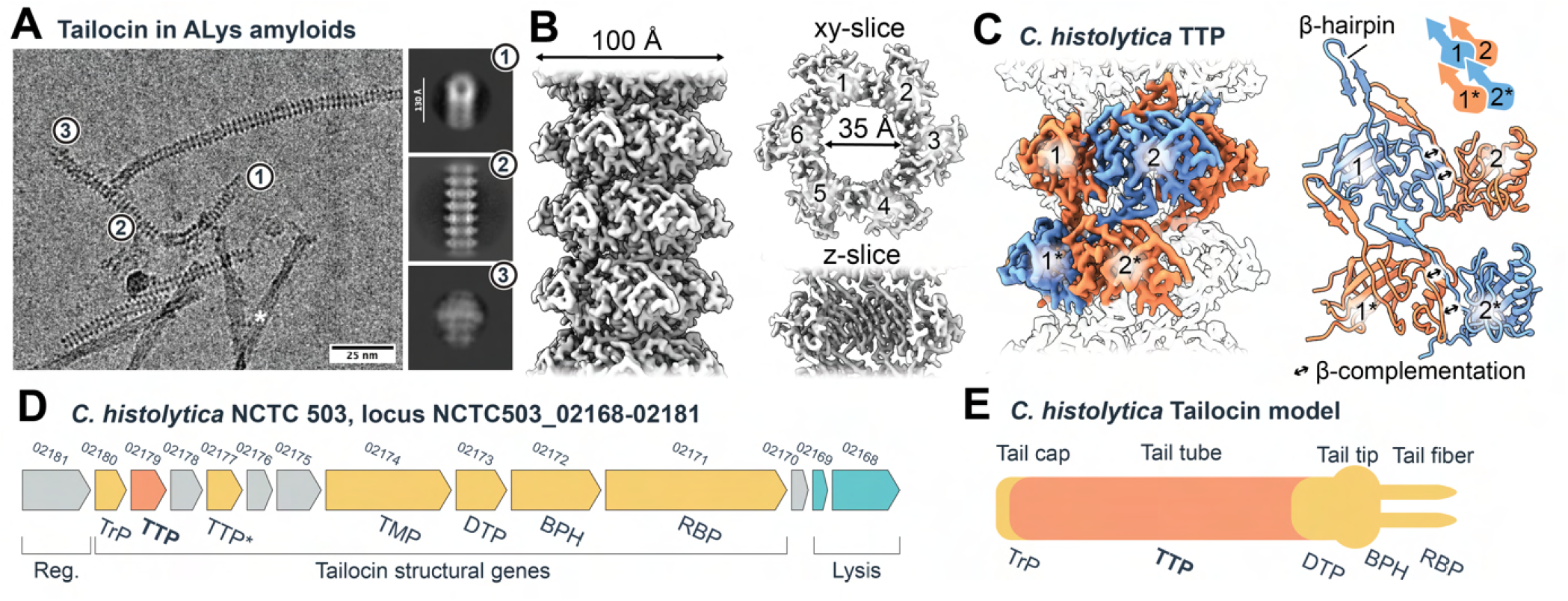
Reconstruction of a major tail tube protein from a *Clostridium histolyticum* tailocin. (**A**) Micrograph from EMPIAR-11785 showing 2D class averages of the tail cap region (1), tail tube (2), and tail tip (3) of a tailocin. Asterisk indicates ALys amyloid. Scale bars: micrograph 25 nm, 2D class 130 Å. (**B**) 3.5 Å helical reconstruction of 10 nm C6-symmetric tubes with a 3.5 nm inner diameter, displayed along the helical axis and as xy- and z-slices. Numerals indicate individual protofilaments. (**C**) Cryo-EM reconstruction color-coded by subunit packing. Inter-protofilament contacts through β-complementation are indicated by black double arrows. **(D)** Schematic representation of a *C. histolyticum* NCTC 503 tailocin element. Grey elements are of unassigned function; yellow indicates structural genes, and genes involved in lysis are in mint. The tail tube protein (TTP) is highlighted in orange. A gene duplicate of the TTP is annotated as TTP*. Tail terminator TrP, tape measure protein TMP, baseplate hub BPH, and tail tip/side fibers RBP are annotated. **(E)** Schematic representation of a tailocin particle with the TTP highlighted in orange.

To identify the assembly, we used automated model building with CryoAtom ^43^, with the human proteome (UP000005640) as a reference database. No convincing matches were identified, arguing against our initial interpretation that the fibrils represented stacks of common amyloid-associated proteins, such as serum amyloid P component (SAP), or intermediates of lysozyme amyloid formation. We then used the fitted poly-alanine backbone trace from automated model building as input for FoldSeek ^45^ to identify proteins with similar folds, which indicated the presence of phage tail tube proteins (TTPs) with *Siphoviridae*-like morphology. Because the global resolution was limited to approximately 3.5 Å, the structure-to-sequence assignment remains probabilistic and relies in part on alignment of aromatic residues (**Fig. S2E**), but the high map-model fit, assessed with the Q-score, supports our assignment (**Fig. S2D**).

To test collagenase as the source of contamination, we performed a mock fibril extraction of the commercial *C. histolyticum* collagenase used for ALys (EMPIAR-11785) and light-chain amyloid extraction. We assumed a reasonably consistent composition of the bacterial secretion used for the manufacturing of collagenase across batches. Our enrichment was designed to retain higher-molecular-weight assemblies with sedimentation properties similar to amyloid fibrils. Subsequent negative-stain transmission electron microscopy (nsTEM) of the enriched material revealed abundant phage tail-like assemblies that resembled the non-amyloid fibrils observed in the Alys dataset. Further processing of the nsTEM data yielded 2D class averages similar to those observed in our cryo-EM data (**Fig. 2A**). Power-spectrum analysis of the prominent layer line revealed a helical rise of about 34 Å, close to the 37 Å rise from our previous cryo-EM reconstruction (**Fig. 2D**). Together, these results indicate collagenase as the reagent-based source of the ALys contamination.

**Figure 2.**
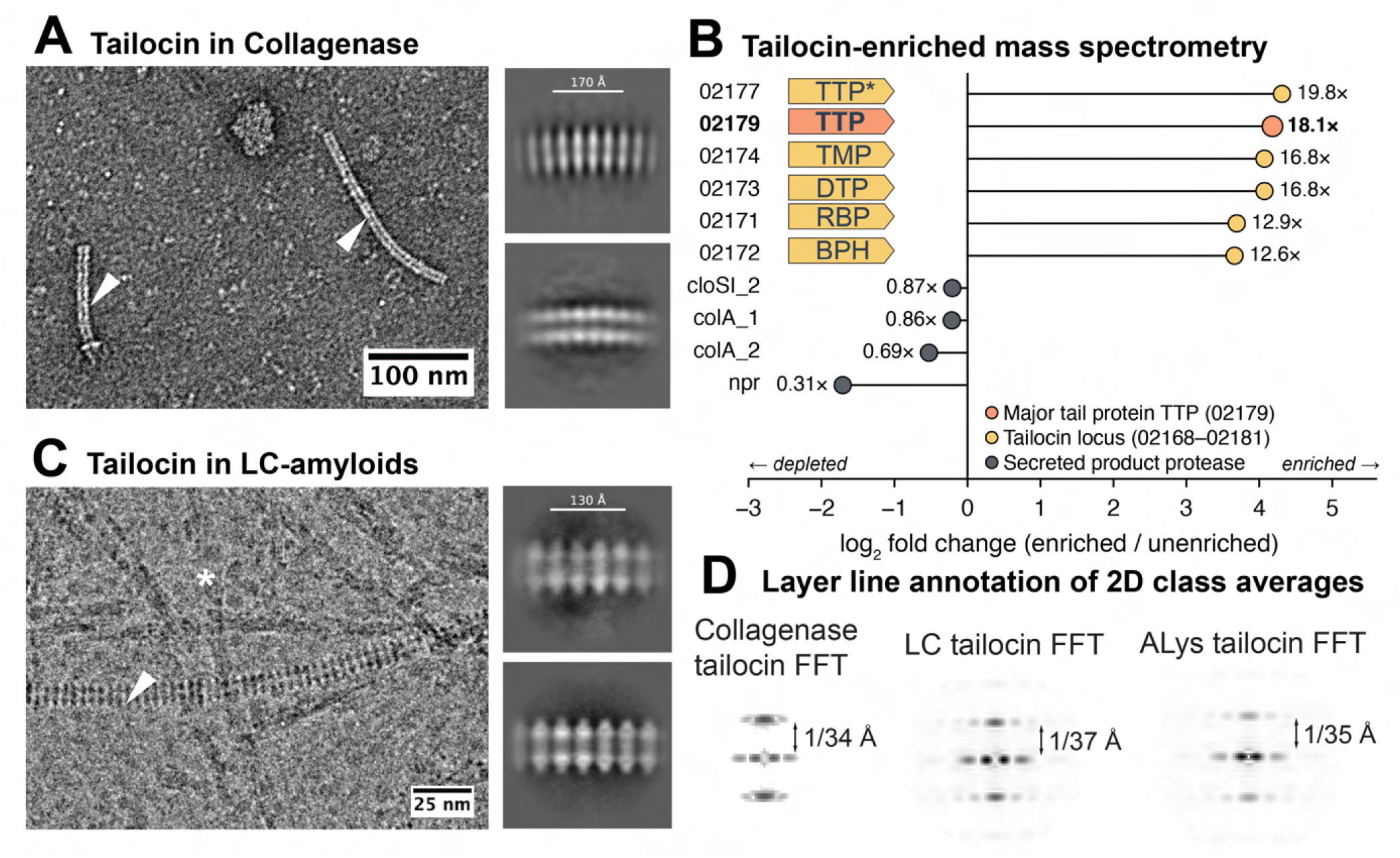
Tailocins are a recurrent contaminant in collagenase-treated ex vivo amyloid extracts. **(A)** Negative-stain TEM micrograph and 2D class averages of tailocin (white arrow) from crude collagenase (Sigma-Aldrich C2674). Scale bars: 100 nm on micrograph, 170 on 2D class. **(B)** log_2_ fold change (limma, two replicates per condition) for all tailocin locus proteins (yellow), the major tail tube protein (orange) and secreted product-protease proteins, and real fold-change as values. Structural tailocin genes annotated according to **Fig. 1D**. (**C**) Micrograph of light-chain amyloids (asterisk) and tailocin (arrow). Scale bar 25 nm. 2D class averages of phage tail fibrils, scale bar 130 Å. (**D**) Comparison of contrast-inverted power spectra of the nsTEM collagenase tailocin (left), cryo-EM of light-chain tailocin (middle panel) and ALys-derived tailocin (right) 2D class averages using the *relion display* function in RELION-5^6^. The annotated layer-line approximates the helical rise of the helical assembly.

To identify the present contaminant, we constructed a candidate sequence database from the *C. histolyticum* proteome (**Table S2**) and inspected the corresponding predicted structures using AlphaFold3 (**Fig. S3**). Subsequently, we performed automatic model building with CryoAtom and the three available *C. histolyticum* proteomes as input. Resulting models were used for another round of FoldSeek. Manual inspection of the FoldSeek targets with the highest confidence scores (**Fig. S3**) supported assignment of the structure to a *C. histolyticum* major tail tube protein MttP, hereafter called TTP, which is part of an F-type non-contractile tailocin element (UniProt: A0A4U9RSA4, **Fig. 1D**).

The same enriched sample and a control were subjected to tryptic digestion and mass spectrometry. Bottom-up LC-MS/MS identified 60 proteins from *C. histolyticum*. Common contaminants were excluded. We detected six proteins encoded in the F-type tailocin locus NCTC503 02168–02181 Å (**Fig. 1D-E**), and all were enriched after mock purification of amyloids. The major tail protein TTP (A0A4U9RSA4), which we characterized structurally, was enriched 18-fold (log_2_ fold change 4.18, adjusted p = 0.028), with 69% sequence coverage by MS/MS from 20 peptides (**Fig. S4**). The subunits assigned by inferred homology with monocin (Lee et al., 2016; Sigal et al., 2024; Gu et al., 2025) to the side fiber / receptor-binding protein (02171), the baseplate hub (02172), the distal tail protein (02173) and the tape measure protein (02174) were enriched 13-to 17-fold, reaching significance for 02171, 02173 and 02174 (adjusted *p* = 0.034, 0.0073 and 0.0091); the comparable enrichment of 02172 (13-fold) did not (adjusted *p* = 0.22). Locus 02177 showed the largest point estimate of the cluster (20-fold) but likewise did not reach significance (adjusted *p* = 0.21); the quantification of 02177 and of 02174 depended in part on imputed values (**Fig. 2B**). The six subunits of a single contiguous locus move together over a narrow 13-to 20-fold range while spanning an approximately 30-fold range in absolute abundance, consistent with their co-enrichment as components of one particle. Rather than the assembly chaperone suggested by its position in the monocin operon, 02177 encodes a second tail-tube copy (TTP*) that shares 88.1 % sequence identity with locus 02179 (TTP) **Fig. S4A**. Local map-model fit analysis of non-conserved positions using the Q-score supports locus 02179, rather than 02177, as main TTP in the cryo-EM reconstruction (**Fig. S4A**).

Tailocins are phage tail-like bacteriocins and act like molecular needles, evolved to puncture membranes of competitor cells under nutrient-scarce conditions. Bacteria encode one or rarely two gene clusters for tailocins. Lytic release of tailocins occurs in a small fraction of cells, gaining overall population fitness. These capsid-free phage tail-like assemblies consist of a tail tube and baseplate with fibers selective for cell surface markers like the O-antigens of lipopolysaccharides ^14,31,38^. Since the observed structure lacks a sheath, it can be categorized as a non-contractile F-type tailocin, structurally related to tails of phages of the *Siphoviridae* family ^3^. Limited by the number of tailocins, we were only able to reconstruct the TTP as part of the structural gene module (**Fig. 1D**). While visible at the micrograph level and in 2D class averages, the tail cap (**Fig. 1A**, panel 1) and tail tip (**Fig. 1A**, panel 3) were not reconstructed. Each protomer within the hexameric TTP stack contains a conserved β-sandwich domain that lines the central channel and an extended β-hairpin involved in inter-stack interactions (**Fig. 1C**). Overall, we identified a commercial enzyme widely used in amyloid research as a source of contamination with bacterial tailocin particles.

### *Clostridium histolyticum* tailocin is a recurrent contaminant in amyloid extracts

We next asked whether tailocins are a recurrent contaminant in ex vivo amyloids. To address this, we analyzed cardiac light-chain amyloids (AL), which cause systemic amyloidosis and arise from the overproduction of immunoglobulin light-chain fragments. Clinically, AL amyloidosis can affect the heart, digestive tract, kidneys, nervous system, and soft tissue ^1^.

Following an established collagenase-based extraction protocol, we enriched light-chain amyloids from the apex of a heart explanted from an AL amyloidosis patient for cryo-EM. Using reference-based particle picking, we also identified tailocin in the AL dataset (**Fig. 2C**). While nearly one-third of ALys micrographs contained observable tailocins, only approximately 5% of our AL micrographs contained the pseudo-phages (**Fig. 2C**). These percentages were estimated by linking particles assigned to tailocin 2D class averages back to their original micrographs. Because of the limited number of particles in the AL dataset, we were unable to reconstruct the AL-associated major tail-tube protein. Using the 2D class averages of the proposed tailocin to generate an average power spectrum in Fourier space, we estimated a helical rise of approximately 34 Å for collagenase tailocins (**Fig. 2D**, left panel). The same approach yielded a rise of about 35 Å for ALys-derived tailocin and approximately 37 Å for light-chain-derived tailocin (**Fig. 2D**, right and middle panel, respectively), further supporting our assessment of tailocin contamination from the use of crude *C. histolyticum* secretion in ex vivo amyloid extracts.

### Strategies to avoid artefacts in cryo-EM

Modern SPA cryo-EM data processing enables reconstruction of minimally processed samples, in many cases omitting affinity-based purification steps entirely. This capability can reveal transient or unstable complexes, but minimally invasive extraction methods also increase the risk that reagent- or preparation-derived artefacts will be overinterpreted as biological findings. Even when sample heterogeneity is deliberately explored, unexpected results should be carefully assessed within the context of sample preparation before they are interpreted as novel biological assemblies. The following set of strategies may help the cryo-EM community identify contaminants, avoiding mis- or overrepresentation of new structures:

#### Reagent-only controls

Mock-protocols or extraction blanks are widely used in sequence-based fields ^34^, as they rely on low-input samples that carry a high risk of reagent-derived signals being misinterpreted as novel biology ^11,28^. Current SPA cryo-EM pipelines lack an equivalent but would benefit from extraction blanks, implemented in nsTEM prescreening. Additionally, testing lot-level reagent quality is advisable, as production quality, especially for biologics, can vary.

#### Genetically encoded contaminants

Host-derived proteins can mimic affinity tags, competing with the intended target. Documented examples are *E. coli* ArnA and AcrB as enriched contaminants in immobilized metal affinity chromatography ^4,7^. Alternative purification routes, such as ion-exchange chromatography or tandem affinity purification, have proven effective amongst several other strategies.

#### Shared properties enrich contaminants

Enrichment strategies involving detergent insolubility ^20^, protease resistance ^5^, or a high sedimentation coefficient ^22,26,27^ will co-purify abundant macromolecules with shared properties of the intended target molecule. Consequently, a protocol selective for physical properties should be described as enriching a class of particles, not a single target.

#### Sequence-agnostic model building

Sequence-agnostic backbone tracing followed by map-based sequence identification allows an unbiased map-model fit from a defined candidate library, covering the proteome of the expression host and intended target sequence ^18,43,45^. This approach is especially valuable for cases in which the composition of the sample is unknown in advance or for unexpected contaminants or co-binders of the intended target ^15,23,40^. The ranked candidate list should be reported in full, including candidates of high backbone but low per-residue map-to-model fit. In cases of limited resolution, probabilistic sequence assignment and fold searches with FoldSeek ^45^ can be explored. For the reported tail-tube fold, which is shared across siphoviral tails and their bacteriocin derivatives ^47^, a single fold assignment can implicate an entire class of contaminants.

#### Annotate deposited maps as contaminants, toward a contamination database for cryo-EM

Contaminant reconstructions are often discarded, causing laboratories to unknowingly rediscover them independently. Proteomics addressed this problem with a shared contaminant database, and crystallography with ContaMiner and ContaBase ^16^. However, cryo-EM has no equivalent resource. Depositing identified contaminant maps to the EMDB with an explicit reagent-derived annotation, as has been done for a phage tail traced to a contaminated buffer ^19^ (PDB 9MFE), would convert the experience of individual research groups into a searchable community asset, allowing depositors to check new maps against known artefacts as a routine step.

## Material & Methods

### Amyloid light-chain extraction

This study, conducted with anonymized ex vivo human cardiac tissue samples obtained by a generous gift from Protego Bio from a donor who provided informed consent, was approved by the Institutional Review Board of Scripps Research (protocol No.: IRB-24-8382, continuing IRB-16-6864 and IRB-12-5994). Light-chain amyloids were extracted from human heart tissue using a water-based protocol ^**?**^. Briefly, 125-250 mg of cardiac tissue from the apex was cut, washed three times with 0.5 mL Tris-Calcium buffer, and incubated overnight with *C. histolyticum* collagenase (0.5 mg/mL) at 37 °C. The sample was pelleted at 3100 x g for 30 min at 4 °C, and the supernatant was discarded. The pellet was resuspended in 0.5 mL Tris-EDTA, centrifuged, and the supernatant was discarded. After 10 total extraction cycles, the pellet was resuspended in 0.5 mL ice-cold water and stored at 4 °C.

### Negative staining TEM of *Clostridium histolyticum* collagenase

Clostridium histolyticum collagenase (Sigma Aldrich, C2674) was resuspended in TBS buffer (50 mM Tris-HCl, 5 mM CaCl_2_) at a final concentration of 4 mg/ml. To enrich higher-molecular-weight components, 0.8 ml sample was sedimented for 30 min at 200,000 x g; the pellet was washed with TBS buffer, resuspended in 80 µl TBS, and assessed by negative-stain electron microscopy following established protocols ^9^. In brief, 100x and undiluted samples were applied to glow-discharged 400 square Mesh Copper/Rhodium Maxaform grids (Electron Microscopy Sciences) and stained with 2 % uranyl formate. Micrographs were collected automatically with Leginon ^44^ on a Thermo Fisher Talos F200C transmission electron microscope operating at 200 keV and equipped with a Gatan K2 direct electron detector at a nominal magnification of 73,000, corresponding to a pixel size of 1.981 Å. Micrographs were imported in cryoSPARC (v.4.7) with constant CTFs and inverted contrast. Templates were generated from manually picked particles. 2D classification without Force max over poses/shifts yielded well-aligned 2D averages.

### Comparative power spectrum analysis

2D class averages of ALys, AL, as well as negative-stain data of *Clostridium histolyticum* collagenase non-amyloid fibrils were opened with the *relion display* function in RELION-5^6^ and converted to their power spectrum. The first layer line distance was manually measured from the equator and converted to the helical rise value according to established protocols ^35^, using: rise [Å] = (box size [px] / FFT layer line distance) * 2 * pixel size [Å /px].

### Mass-spectrometric characterization of *Clostridium histolyticum* collagenase

#### MS Sample preparation

Collagenase and mock enriched collagenase were prepared for bottom-up proteomic analysis by denaturation in TBS containing 2% (w/v) SDS. Cysteine residues were reduced with 5 mM tris(2-carboxyethyl)phosphine (TCEP) for 30 min at room temperature and subsequently alkylated with 10 mM iodoacetamide (IAA) for 30 min at room temperature in the dark. Proteins were digested overnight at 37°C with trypsin at an enzyme-to-protein ratio of 1:100 (w/w). Digestion was terminated by addition of trifluoroacetic acid (TFA) to a final concentration of 2%, followed by incubation for 5 min at 37°C with shaking at 800 rpm. Peptides were subsequently desalted using C18-based cleanup, dried by vacuum centrifugation, and reconstituted in 0.1% formic acid prior to LC–MS/MS analysis.

#### LC–MS/MS analysis

Peptides were analyzed by nanoLC–MS/MS using a Vanquish Neo UHPLC system coupled to an Orbitrap Exploris 480 mass spectrometer (Thermo Fisher Scientific) operated in data-dependent acquisition (DDA) mode. Peptides were loaded using a trap-and-elute configuration with a 300-µm × 5-mm trap column and separated on a 75-µm × 50-cm analytical column at a flow rate of 350 nL/min. Mobile phase A consisted of 0.1% FA (Formic acid) and mobile phase B consisted of 80% acetonitrile. The gradient began at 2% B, increased to 25% B over 75.1 min and to 38% B by 112.1 min, followed by a column wash at 99% B. The total method duration was 120 min. Peptides were ionized by nanospray ionization in positive-ion mode at 1.9 kV. MS1 spectra were acquired in the Orbitrap at a resolution of 120,000 over an *m/z* range of 320–1,600 with a normalized AGC target of 300%. Precursors with charge states of 2–6 were selected for fragmentation using a 3-s cycle-time method, with dynamic exclusion of previously selected precursors for 30 s. MS/MS spectra were acquired following higher-energy collisional dissociation (HCD) using a normalized collision energy of 28%, a precursor isolation window of 1.6 *m/z*, an Orbitrap resolution of 15,000, a normalized AGC target of 100%, and a maximum injection time of 50 ms.

#### Database search and identification of *C. histolyticum*-derived peptides

Raw LC–MS/MS data were processed using FragPipe v24.0 with MSFragger v4.4.1. Spectra were searched against a target-decoy database containing the relevant *C. histolyticum* proteome (UniProt proteome UP000308489), together with common contaminant sequences. Trypsin specificity was specified with up to two missed cleavages, and peptides between 7 and 50 amino acids were considered. Precursor and fragment mass tolerances were set to ±20 ppm and 20 ppm, respectively. Carbamidomethylation of cysteine (+57.02146 Da) was specified as a fixed modification, while methionine oxidation (+15.9949 Da) and protein N-terminal acetylation (+42.0106 Da) were included as variable modifications. Database-search results were rescored using Percolator v3.7.1, followed by protein inference using ProteinProphet. Protein identifications were filtered using a target-decoy strategy with a 1% protein-level false discovery rate. Label-free peptide/protein quantification was performed using IonQuant v1.11.20 with match-between-runs enabled.

### Cryo-electron microscopy of cardiac light-chain amyloids

UltraAuFoil 200 mesh R 2/2 grids were glow-discharged under vacuum for 30 s at 15 mA in a Pelco 311 easiGlow 91000 glow discharge cleaning system (Ted Pella). 3.5 µl sample was applied to the front of the grid and 0.5 µl to the back, incubated for 1 min, and backblotted with Whatman 1 filter paper for 4-6 s after the liquid spot on the filter paper stopped spreading. Grids were manually plunge-frozen in a 4 °C cold room at *>*95% humidity. AutoGrids were assembled and immediately screened.

Cryo-EM data were collected on a Talos Arctica TEM (Thermo Fisher) operating at 200 keV. Movies were recorded using a Falcon 4i direct electron detector (Thermo Fisher) at a nominal magnification of 150,000, corresponding to a pixel size of 0.94 Å. Movies were saved in the electron-event representation (EER) format and recorded with a total electron exposure of 50 e^-^ per Å ^2^. All datasets were collected automatically using EPU (v.3.9, Thermo Fisher) with a defocus range of -0.8 to -2.0 µm. EPU’s fast exposure navigation was used to collect data with an 8 µm image shift. All datasets were processed using cryoSPARC (v.4.6).

For preprocessing, movies were dose-fractionated into 40 frames and motion corrected using patch-based motion correction, followed by patch-based CTF estimation in cryoSPARC Live. Micrographs with CTF fits worse than 6 Å and astigmatism greater than 600 Å were discarded. The remaining accepted micrographs were used for further processing. Per-dataset micrograph statistics are compiled in **Table S1**.

### Processing of tubular filaments in EMPIAR-11785

From the 2,013 processed micrographs in EMPIAR-11785, approximately 30% contained non-amyloid segments and were selected for further processing. To increase the number of particles, filaments were traced using a diameter range of 40-80 without templates and with 0.5 x Gaussian blur. Segments were extracted in a 208-pixel box, Fourier-cropped to 100 pixels, and subjected to two rounds of 2D classification with an initial class uncertainty factor of 4 and hard classification during the final iteration. A total of 30,000 selected segments were re-extracted with recentering in a 256-pixel box without binning. Helical refinement was performed without symmetry using a cylindrical model and a 60° tilt search-range. The resulting reconstruction was used as input for real-space helical indexing in HI3D ^**?**^ with default parameters, suggesting a twist of 13.74° and a rise of 37.93 Å. These values were used in helix refine with non-uniform refinement (NUR) enabled. To test for higher-order symmetry, Phenix’s symmetry search tool *phenix*.*map symmetry* ^**?**^ was used and identified the highest correlation with C6 point-group symmetry. After another round of 2D classification, 21,000 segments were used in helical refinement with C6 symmetry, a rise of 38 and a twist of 13.7°. Helical indexing with the new C6 point group and higher-resolution map suggested a rise of 37.19 and twist of 14.21°. The final helical refinement with a 20° tilt search-range resulted in a 3.5 Å reconstruction (FSC 0.143). The map was sharpened automatically within cryoSPARC using a B-factor of –56 Å ^2^. The full processing workflow and validation are shown in **Fig. S1, Fig. S2** and **Table S1**.

### Construction of a candidate sequence database

Sequence candidates were assembled from the *Clostridium histolyticum* reference proteome (UniProt UP000308489) carrying phage-, tail-, capsid-, portal-, terminase-, holin- or bacteriocin-related annotations. The corresponding models were retrieved from the AlphaFold Protein Structure Database, with a list of annotated non-redundant candidates provided in supplementary **Table S2**.

### Sequence-agnostic identification of the fibrillar species

A main-chain model was built de novo into the sharpened map using CryoAtom v.2.1.0^43^ in no-sequence mode, and the resulting per-residue amino-acid probability profiles were used for hidden Markov model searches (*hmm search*) against three databases in turn: (I) the 22-protein candidate set; (II) the three available *Clostridium histolyticum* proteomes (UP000308489, UP001732247 and UP001732502); and the human reference proteome (UP000005640). In parallel, AlphaFold Protein Structure Database models for all 22 candidates were rigid-body docked into a single asymmetric unit of the map in ChimeraX v.1.10.1 and ranked by the map-model fit Q-score plugin. Structures shown in **Fig. S3**.

### Model Building and Refinement

Map resolvability was improved through postprocessing in Å EMready2^8^. The AlphaFold model of A0A4U9RSA4 was manually built into the density in Coot v.0.9.8.95^12^ and ISOLDE ^10^. The helical assembly was generated by applying the refined helical parameters with the *sym* command in ChimeraX, and the resulting dodecamer model was refined against the sharpened map in *phenix*.*real space refine* ^25^ with secondary-structure, Ramachandran and non-crystallographic symmetry restraints. Further validation was performed using the Q-score ^33^ for residue-wise quality estimation. The best model was further refined using ISOLDE and Phenix. Results were visualized using ChimeraX v1.11.1^32^.

## Data availability

All re-processed datasets from the Electron Microscopy Public Image Archive (EMPIAR) were deposited in the Electron Microscopy Data Bank (EMDB) and Protein Data Bank (PDB). The datasets from Table S1 are available using the following Å EMDB accession numbers with their respective EMPIAR-ID: *Clostridium histolyticum* TTP (PDB-11OK, EMD-72138, Å EMPIAR-11785). Mass spectrometry data are available via ProteomeXchange with identifier PXD084014.

## Acknowledgments

Funding for this research was provided by a NINDS grant NS095892 to GCL, a NIDDK grant DK046335 to JWK, and the German Research Council project number 556478029 to JHS. RTO is grateful for the financial support from the George E. Hewitt Foundation for Medical Research. The authors thank Jean-Christophe Ducom and Charles Bowman at Scripps for computational support. BioRender was used to create parts of the graphical abstract.

## Declaration of generative AI and

### AI-assisted technologies in the manuscript preparation process

During the preparation of this work, the authors used Claude to assist in writing code for data extraction and visualization. After using this tool/service, the authors reviewed and edited the content as needed and take full responsibility for the content of the published article.

### CRediT authorship contribution statement

Jan-Hannes Schaefer: Conceptualization, Data curation, Formal analysis, Funding acquisition, Investigation, Methodology, Resources, Software, Validation, Visualization, Writing – original draft. Zi Gao: Investigation, Methodology, Writing – review and editing. Robert T. O’Neill: Investigation, Writing – review and editing. Danielle A. Grotjahn: Supervision, Writing – review and editing. Evan T. Powers: Supervision, Writing – review and editing. Jeffery W. Kelly: Funding acquisition, Project administration, Supervision, Writing – review and editing. Gabriel C. Lander: Funding acquisition, Project administration, Supervision, Writing – review and editing.

## Supporting Information

**Figure S1.**
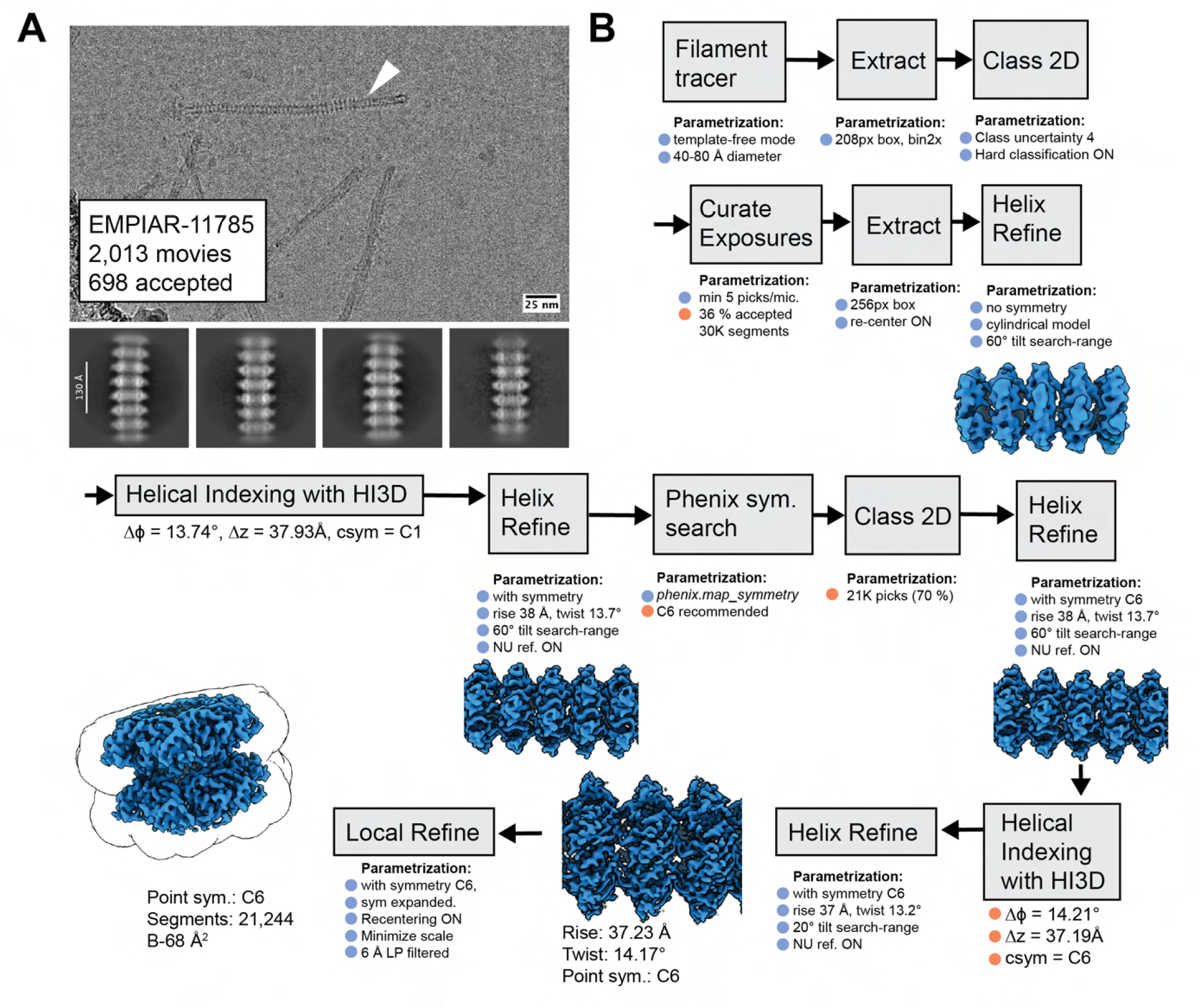
Reconstruction of a *C. histolyticum* tailocin tail tube protein. **A** Micrograph and 2D class averages from EMPIAR-11785 positive for the pseudo-phage tails (white arrow). 25 nm scale bar on micrograph and 130 Å scale bar on 2D class averages. **B** Processing workflow for helical reconstruction.

**Figure S2.**
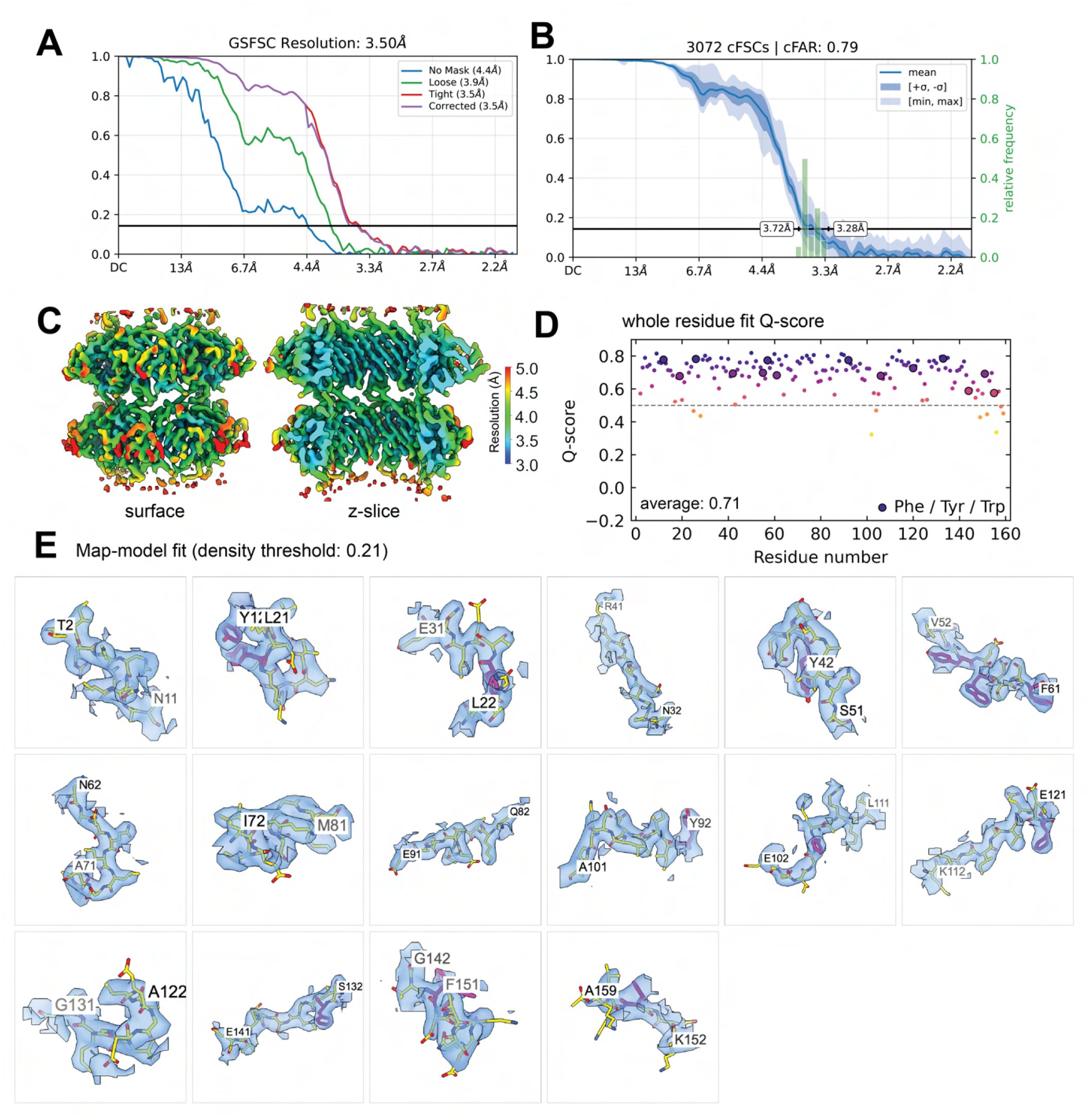
Validation of a *C. histolyticum* tailocin tail tube protein. **A** Gold-Standard FSC resolution at threshold 0.143. **B** Orientation diagnostics, including Conical FSCs (cFSC) and Conical FSC area ratio plots (cFAR). cFAR ¡ 0.5 indicates anisotropic reconstruction. **C** Local resolution estimates with surface- and z-slice representation.**D** Atom resolvability analysis using the whole-residue Q-score plot and a global average of 0.71. Aromatic residues highlighted and Q-score 0.5 indicated with a dashed line. **E** Map-model fit of the entire monomeric TTP in tiles of 10 amino acid stretches at a density threshold of 0.21. Aromatic residues highlighted in violet.

**Figure S3.**
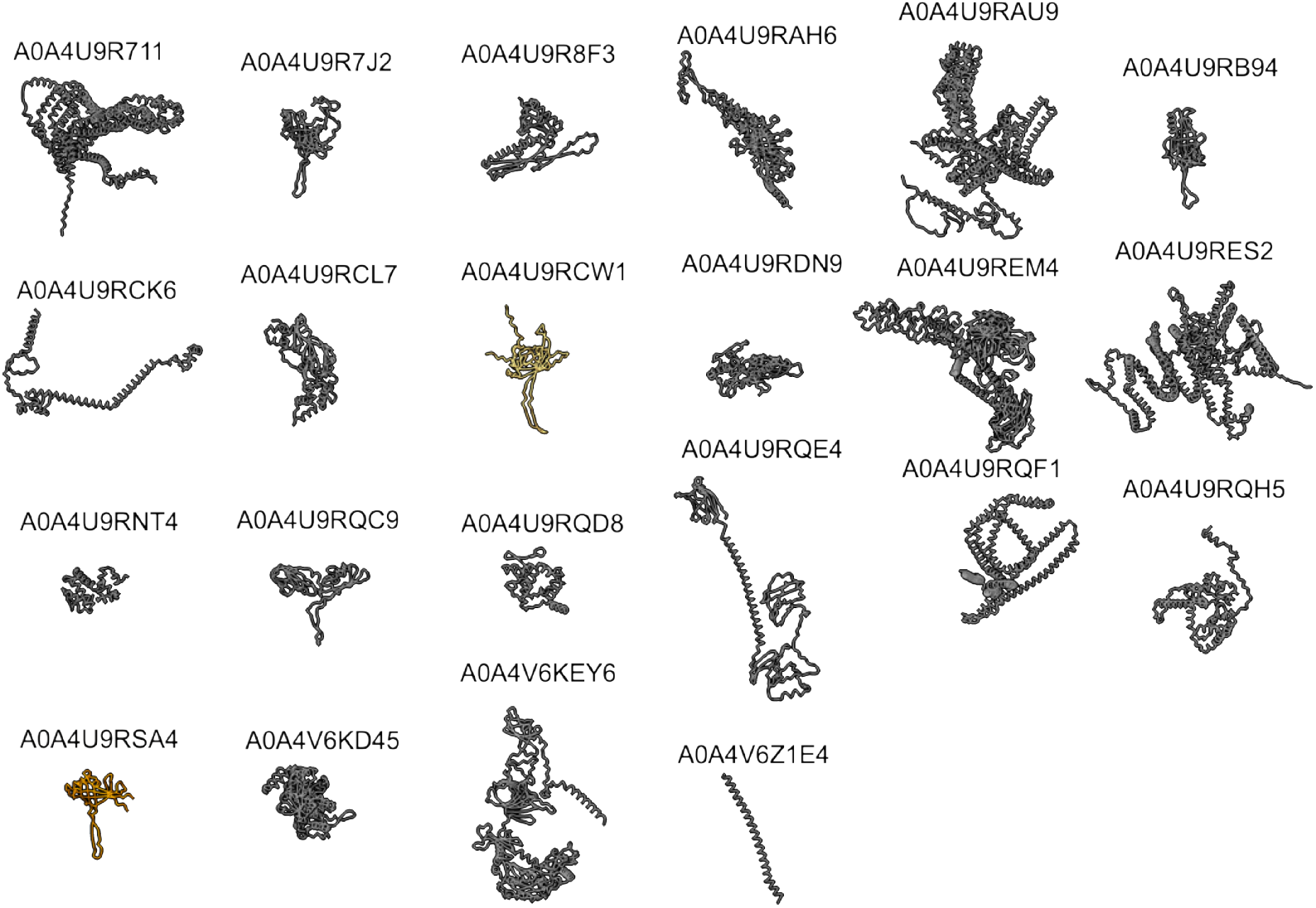
Library of candidate structures. AlphaFold models (from a recent server API call) of candidates identified from the *C. histolyticum* genome in Table S2. Structures in gray show poor model-map fit, whereas candidates in yellow yield better model-map fits with rigid docking in ChimeraX. The best q-score was calculated for model A0A4U9RSA4, *C. histolyticum* locus 02179 in orange.

**Figure S4.**
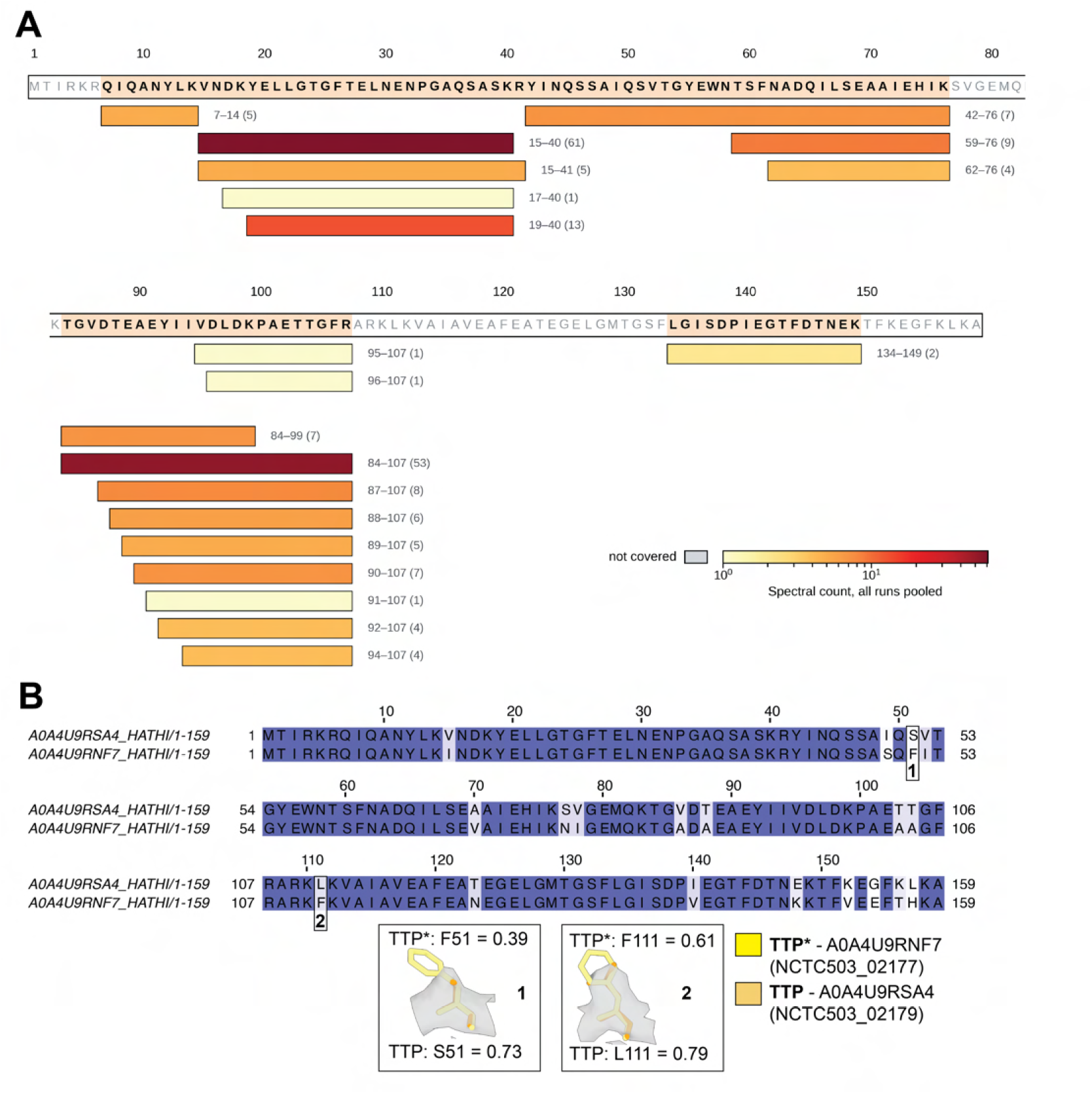
**A** MS/MS peptide sequence matches. A0A4U9RSA4 phage major tail protein (NCTC503 02179) – 159 aa, 69 % sequence coverage, 204 assigned spectra (all runs pooled). **B** Paired sequence alignment of tail protein loci in the tailocin element of *C. histolyticum* colored by sequence identity. Inserts 1 and 2 show map-to model fits of diverging sequence identity using Q-score values at position 51 (insert 1) and position 111 (insert 2).

**Table S1.** Cryo-EM data collection, image processing, and modeling statistics of *C. histolyticum* major tail tube protein (TTP).

|  | **TTP** |
| --- | --- |
| EMDB ID | 72138 |
| EMPIAR ID | 11785 |
| PDB ID | 11OK |
| Microscope | Krios |
| Camera | K2 Quantum |
| Magnification (nom.) | 130,000 |
| Voltage (keV) | 300 |
| Total dose (e<sup>-</sup>/Å<sup>2</sup>) | 49 |
| Exposure rate (e<sup>-</sup>/px/s) | 5.9 |
| Frames per movie | 40 |
| EER Fractions | NA |
| Pixel size (Å/px) | 1.04 |
| Defocus range (μm) | −1.0 to −2.0 |
| Recorded movies | 2,013 |
| Processing software | CS (v.4.7) |
| Final particle images | 21,244 |
| Symmetry imposed | C<sub>6</sub> |
| Helical twist (°) | +14.17 |
| Helical rise (Å) | 37.23 |
| Res. (FSC 0.143, Å) | 3.5 |
| Local res. range (Å) | 3.3–3.7 |
| cFAR | 0.79 |
| Sharpening B-factor (Å<sup>2</sup>) | −68 |
| Initial model | AlphaFold (A0A4U9RSA4 ) |
| Refinement | Coot/ISOLDE/Phenix |
| FSCmap-to-model(0.5) (Å) | 3.8 |
| MolProbity score | 1.32 |
| Clash Score | 3.42 |
| Composition |  |
| Chains | 12 |
| Atoms | 29,232 |
| Protein residues | 1,896 |
| Bonds (R.M.S.D.) |  |
| Length (Å) | 0.003 |
| Angles (°) | 0.573 |
| B-factors (min/max/mean) |  |
| Protein residues | 61.69/101.28/74.84 |
| Ramachandran plot (%) |  |
| Favored | 96.85 |
| Allowed | 3.15 |
| Outliers | 0.00 |
| Rotamer outliers (%) | 0.70 |
| Q-score | 0.71 |
| EMRinger score | 3.93 |

**Table S2:**
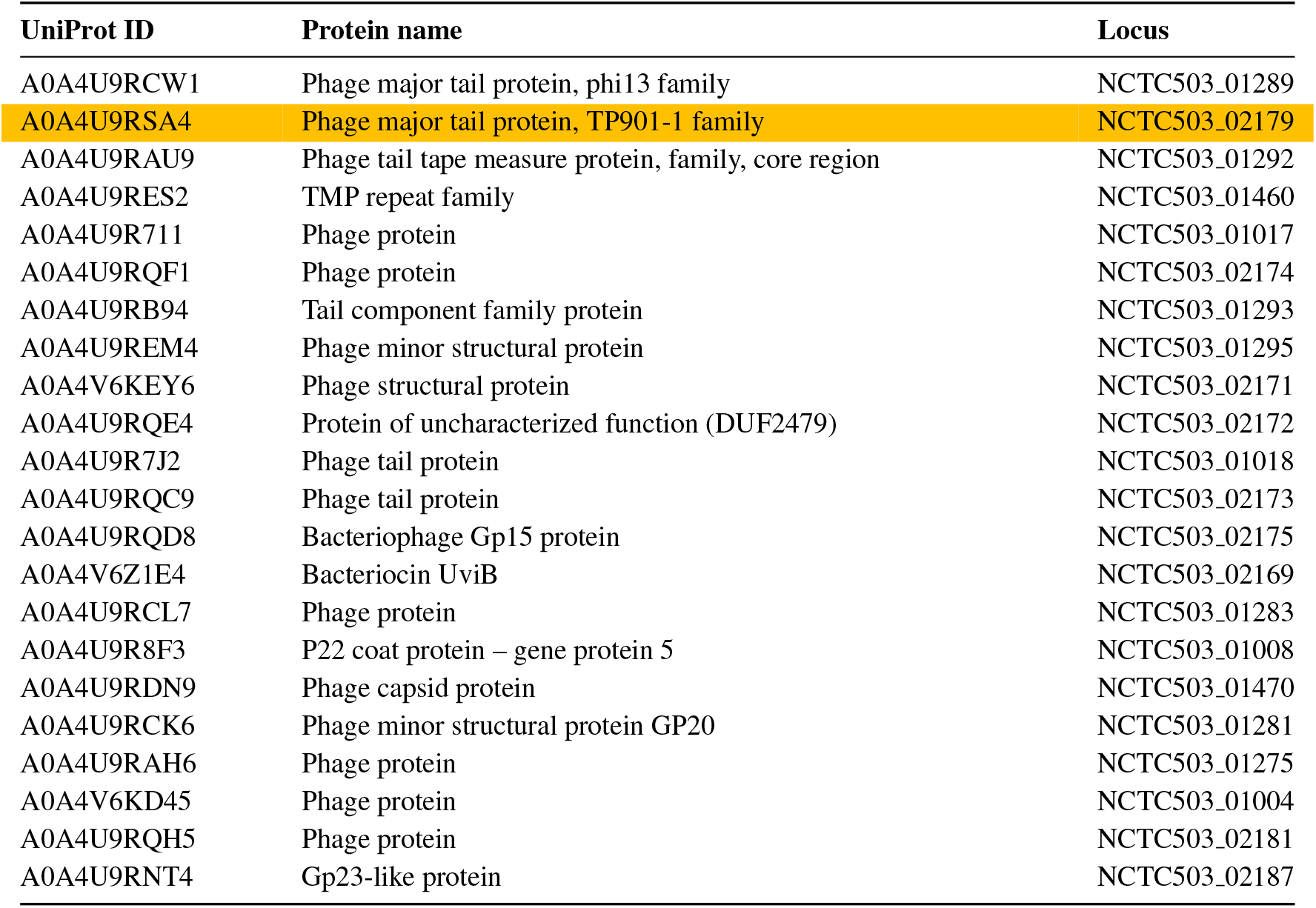
Candidate Library of *C. histolyticum* proteome UP000308489. Final candidate highlighted in orange.

